# Large-Scale Multigenome-Wide Study Predicts the Existence of Transmembrane Phosphotransfer Proteins in Plant MSP Signaling Pathway

**DOI:** 10.1101/2025.07.28.667123

**Authors:** Sergey N. Lomin, Wolfram G. Brenner, Ekaterina M. Savelieva, Dmitry V. Arkhipov, Georgy A. Romanov

## Abstract

A new class of plant phosphotransfer proteins belonging to multistep phosphorelay (MSP) system was discovered by using large-scale bioinformatics methods. Unlike the canonical soluble nucleo- cytosolic forms, these proteins were predicted to have transmembrane (TM) domains and, apparently, should be localized on some kind of cell membranes. To date, 95 predicted TM-containing phosphotransmitter (TM-HPt) homologs were found in 61 plant species belonging to different clades, taxa and groups: bryophytes, gymnosperms, mono- and dicotyledons. The conserved HPt motif with phosphorylatable histidine was preserved in the most of TM-HPts under study that allowed us to consider these proteins potentially active in MSP signaling. For the found TM-HPts, a Bayesian analysis at the DNA level was performed and a relevant phylogenetic tree was constructed. According to evolutionary relationships, plant TM-HPts were divided into two main groups corresponding to Arabidopsis AHP1-3,5,6 and AHP4 orthologs. Transcriptomic analysis confirmed the expression of the most of the investigated TM-HPt-encoding genes. Their moderate-to-low overall transcription rate may be a consequence of inducible and/or tissue-specific expression. Using molecular modeling methods, a variety of potential spatial organizations of several such proteins are demonstrated. Possible roles of TM-HPts as modulators of MSP signaling pathway and corresponding putative mechanisms of their action are suggested.

## 1. Introduction

Cytokinins are essential plant hormones that use a multistep phosphorelay (MSP) system for intracellular signal transduction [1–4]. These hormones are involved in diverse processes of plant growth and development such as formation of shoot meristem and leaves, cell division, vascular development, chloroplast differentiation and leaf senescence as well as responses to biotic and abiotic stresses. The cytokinin signaling is triggered by the specific binding of the cytokinin ligand to the receptor and leads to changes in gene expression [5–8].

The cytokinin signal transduction pathway involves a His-Asp phosphorelay similar to that found in bacterial two-component signaling systems [7,9]. But, unlike canonical bacterial signaling which occurs via phosphotransfer between a conserved His residue in the sensor kinase and a conserved Asp residue in the receiver domain of the cognate transcription factor [10,11], in plants this phosphorelay is called MSP, which includes three types of proteins: hybrid receptors with both histidine kinase (HK) and receiver domains, conserved histidine-containing phosphotransfer proteins or phosphotransmitters (HPts) and type B response regulators (RRBs) which are transcription factors containing a conserved phosphoaccepting Asp residue. In the commonly recognized scheme of cytokinin signal transduction, HPts play a special role linking the perception of cytokinins by membrane receptors to the activation of cytokinin-responsive transcription factors, RRBs, in the nucleus. The conserved His residue of the HPts accepts the «hot» phosphate from a conserved Asp residue of the activated cytokinin receptors, carries it into the nucleus and phosphorylates there the conserved Asp residue of the RR protein. Thereafter, phosphorylated type B RRs become able to activate the target genes [3,12–15]. Therefore, HPts have a distinct function in the response to cytokinins in plant cells [16,17]. Suppression of the HPt genes reduced the cytokinin-triggered upregulation of sensitive genes including type A RRs [18]. Insertional mutations in *AHPs* reduced the sensitivity of plants to the hormone in various cytokinin bioassays [19].

HPts are represented in plants by a family of small proteins with predicted histidine phosphotransfer activity [20]. Arabidopsis has a corresponding family of six members: AHP1-AHP5 that function as *bona fide* HPts whereas APHP1/AHP6 is a pseudo-HPt (PHP) lacking the conserved phosphoaccepting His residue [21]. AHP1-AHP5 exhibit overlapping functions, mostly acting as positive cytokinin signaling compounds [19,22–24]. The complete blocking of the phosphotransmitter gene expression in a quintuple *ahp1-5* mutant resulted in nonviable Arabidopsis seedlings [25]. By contrast, APHP1/AHP6 acts as a repressor of the cytokinin response pathway [26].

It should be noted that HPts and PHPs are apparently able to interact not only with proteins (sensor HKs, RRs) involved in cytokinin signaling, but also with other hybrid histidine kinases, including, for example, Arabidopsis CKI1 kinase [27] and ETR1 ethylene receptor [28, 29]. Hence, HPts are versatile members of MSP, mediating signal output by various hybrid histidine kinases and input to a variety of RR transcription factors [30]. In addition, the functions of PHPs are not the same across plant species. In rice, a member of the monocot family where PHPs are evidently also negative regulators of cytokinin signaling, disruption of PHP genes results in a subset of phenotypes distinct from ones of the analogous Arabidopsis mutants. This suggests that monocot and dicot PHPs can play non-identical roles in regulating plant growth and development [31]. Thus, some functions of both HPts and PHPs have yet to be uncovered.

In contrast to the remaining uncertainty in the peculiar functions of HPts and PHPs, the question about their localization seemed to have been resolved long ago. Initially HPts (in particular AHPs) were reported to be localized to the cytosol in unstimulated cells and accumulate in the nucleus upon cell exposure to the exogenous cytokinin [16]. However, subsequent quantitative analysis revealed that the subcellular localization of the AHPs is constantly nucleo- cytosolic irrespective of the state of cytokinin signaling. Up to date it is believed that AHPs are soluble proteins constantly cycling between the nucleus and the cytosol [32]. This localization seems logical given the function of HPts in MSP and, particularly, cytokinin signal transmission. Intriguingly, our recent bioinformatics analysis predicts the presence of transmembrane (TM) domains in some HPts. In this work we carried out a global search and analysis of TM-containing plant HPts using bioinformatics methods. As a result, 95 putative plant TM-HPts were found, their domain architecture and biosynthesis at the transcription level were described, and a phylogenetic tree was constructed. Molecular models of a number of these proteins were built. Based on the data obtained, the putative mechanisms of action of TM-HPts were suggested.

## 2. Methods

### 2.1. Bioinformatics Search for Phosphotransmitters with TM Domains

The algorithm of a program specially created for TM-HPt search is shown schematically in the **Figure S1**. Based on the NCBI Protein Reference Sequences database [33], the program performed an automatic search and alignment using NCBI BLAST [33] (protein-protein BLAST algorithm, refseq_ and nr_ databases) of all TM-HPt homologs available in the NCBI database. The retrieved sequences were automatically passed through the «one species - one protein» filter (SINGLESPEC_FILTER, see **Figure S1**). This filter uses the species name of the protein contained in the FASTA sequence metadata. All non-plant proteins were manually removed from the resulting pool of sequences (STAGE-1 OUTPUT). Selected plant sequences were used to repeat the first step (search and alignment of homologs). This time each of the plant-belong sequences from STAGE-1 OUTPUT in turn was used as a query sequence. The number of output sequences was limited to 100,000 proteins. Repeated proteins were automatically removed. The remaining data were automatically screened using the TMHMM 2.0 [35,36] and Phobius [37,38] algorithms for the presence of predicted TM domains.

All proteins for which at least one of the services predicted the presence of TM domains were manually checked using the CCTOP service [39,40] to exclude proteins from the working sample whose detection of a TM domain is an artifact of a specific predictor program. Upon this check, another small part of the proteins was eliminated. Search in the databases Phytozome [41], PlantGenIE [42], Hornworts [43] and Fernbase [44] were performed manually using the BLAST(p) tool. All found HPts passed through the TM filter described above. All selected plant proteins with predicted TM domains were checked on the InterPro website [45,46] for the presence of HPt domain(s). If the HPt domain of a protein was defined by at least one of the databases integrated into InterPro, we kept it in our set.

### 2.2. Phylogenetic Analysis

We have analyzed the nucleotide protein-coding sequences of 64 genes. Each such sequence corresponded to one gene. They were aligned using Clustal Omega [47]. Phylogenetic trees were constructed using MrBayes-3.2.7 [48]. Clustal alignments were used as input, and Bayesian MCMC phylogenetic trees were constructed based on Markov chain Monte Carlo simulation with the general time reversible (GTR) nucleotide substitution model and site-rate variation drawn from a discrete gamma distribution with six classes. 1000000 generation were taken for reaching of standard deviation of split frequencies below 0.01. Resulted tree were visualized by FigTree v1.4.4 [49].

### 2.3. Determination of the Expression of TM-HPt-Encoding Transcripts

To determine whether annotated transcripts encoding TM-HPt proteins were expressed, we employed a large-scale transcriptomic approach. The RNA-sequencing datasets used are listed in Table S1. RNA-sequencing data of large experiments were downloaded from the NCBI SRA database using the prefetch tool of the NCBI SRA Tools suite [50]. The SRA files were extracted to FASTQ files using the fasterq-dump tool. The FASTQ files were quality-checked using the FastQC [51] implementation in Unipro UGENE [52–54].

The reads were mapped to the templates using Rsubread [55]. Templates for mapping the reads were generated as FASTA files containing the annotated mRNA encoding the TM-HPt, or – if not available – the corresponding genomic sequence. In addition, the mRNAs of the orthologs of the *Arabidopsis thaliana* genes At5G53300 (encoding UBC10) and AT3G25800 (encoding PP2AA2) were included as references.

The resulting BAM files were merged using UGENE. The merged BAM files were imported into UGENE to visualize the mapped reads. UGENE was used to determine the number of reads mapped. The maximum number of reads was used as an estimate for expression of the TM-HPt transcripts. If the TM-HPt transcript was a splicing variant, only the TM-HPt-specific part of the transcript was considered. This is marked with “exon” in the tables and figures. To normalize for differing sequencing depths between the species, the number of mapped reads per million total reads was calculated and plotted in the resulting graph. The entire process from downloading the SRA files to mapping with Rsubread was automated in a custom-made R script using the metadata of the SRA dataset as a starting point.

### 2.4. Molecular Modeling

Molecular modeling by *de novo* method was performed using the IntFOLD (version 7.0) web service [56,57]. The best models were selected based on both IntFOLD score and the relevance of protein folding and topology to potential localization in the membrane. The models were optimized in Yasara Structure software (version 22.9.24) [58] using the md_refine macro. This macro runs MD simulations for a 500 ps model using the protocol described in [59]. The pH value was set to 7.4. Prediction of TM regions and signal peptides/sequences was based on Phobius [37,38] implemented in the Protter web service [60,61] which was also used for identification of N-glyco motifs. Orientation of modeled proteins in membrane was predicted in PPM 3.0 Web Server [62,63] and Yasara Structure. Models were additionally optimized and visualized in UCSF Chimera software (version 1.14) [64].

## 3. Results and Discussion

### 3.1. Search and Identification of Potential TM Phosphotransfer Proteins

Previously, while studying and characterizing components of the cytokinin signaling pathway [15,65], we unexpectedly found that protein structure prediction servers identified TM domains in some HPts. Such a TM structure is not only uncharacteristic of plant HPts, but seems to contradict their known function. However, these data raised the issue of the existence of such proteins and their presence in various species. To address this issue, we have undertaken a large-scale search for potential TM-HPts using primarily the available sequenced genomes of higher plants.

The first stage of the search for TM-HPts was carried out in a semi-automatic mode in the NCBI database [33]. A description of the created program and its search algorithm are available in the Section 2.1. This program allowed us to cover all plant species whose genomes were available in this database. The second stage of the search was carried out manually in the databases Phytozome, PlantGenIE, Hornworts, and Fernbase [41–44]. As a result of the search, 95 potential plant HPts with predicted TM domains were found, 82, 10, 2 and 1 in NCBI, Phytozome, PlantGenIE and Hornworts databases, respectively.

Thus, the working set was formed from almost a hundred proteins. The number of TM domains and their positions were determined in each protein using CCTOP and Phobius servers [37,39]. For each selected protein, the potential functionality in MSP was assessed. For this purpose, both the presence of an HPt active site (phosphorylatable histidine) and the putative binding surface (phosphorylation motif) were determined (see **Supplementary Figure S2**). The structure of the conserved phosphorylation motif, defined by aligning sequences of various known active HPts, is shown in **Table 1**. We have specified and extended this motif up to 15 amino acid (aa) residues. The position of the conserved histidine (active site) was set to zero. The positions upstream and downstream of the conserved histidine are indicated by negative and positive numbers, respectively. HPts are considered active if they have phosphorylation motifs that strictly correspond to a conserved aa sequence. When only one aa residue deviated from the perfect motif, we still considered the corresponding proteins to be potentially active. In other cases, the conserved motif was assumed to be not preserved and the respective protein not a functional HPt (**Table 2, Supplementary Table S2**).

**Table 1.** Conserved His phosphorylation motif in active HPts. Sequence of typical aa adjacent to phosphorylatable His residue is indicated, along with their helix and space localization.

| Position aa number | | Typical aa on this position | α-helix | Facing outside or inside |
| --- | --- | --- | --- | --- |
| H set to 0 | in AHP2 |  |  |  |
| -1 | 81 (V) | V, M | $\alpha 4$ | in |
| **0** | **82 (H)** | **H** | $\alpha 4$ | out |
| 1 | 83 (Q) | Q | $\alpha 4$ | out |
| 2 | 84 (L) | L, F, Y | $\alpha 4$ | in |
| 3 | 85 (K) | K | $\alpha 4$ | out |
| 4 | 86 (G) | G | $\alpha 4$ | out |
| 5 | 87 (S) | S | $\alpha 4$ | out |
| 6 | 88 (S) | S | $\alpha 4$ | in |
| 7 | 89 (S) | S, A, T | $\alpha 4$ | out |
| 8 | 90 (S) | S | $\alpha 4$ | out |
| 9 | 91 (V) | V, I | $\alpha 4$ | out |
| 10 | 92 (G) | G | $\alpha 4$ | in |
| 11 | 93 (A) | A | $\alpha 4$ | in |
| 19 | 101 (V) | V, I, A, L, T, S, M | $\alpha 5$ | out |
| 22 | 104 (K) | K, R, Q | $\alpha 5$ | out |

**Table 2.** List of some potential TM-HPts and their main characteristics.

| Plant species | Protein ID | Database | Protein length (aa) | Predicted TM position: |  | Active site | Binding motif |
| --- | --- | --- | --- | --- | --- | --- | --- |
|  |  |  |  | CCTOP | Phobius |  |  |
| *Actinidia eriantha* | XP_057508182.1 | NCBI | 211 | 17-34 | 20-39 | + | + |
| *Anthoceros punctatus* | Apun_evm.model.utg0000231.837.1 | Hornworts | 241 | 198-215 | 199-219 | + | + |
| *Aquilegia coerulea* | Aqcoe7G149700.1.p | Phytozome | 224 | 25-52 | 12-32<br>38-57 | + | + |
| *Arachis hypogaea* | XP_025611698.1 | NCBI | 159 | 14-33<br>52-67 | 51-68 | + | + |
| *Brachypodium arbuscula* | Barbu.8G252800.1.p | Phytozome | 153 | 132-150 | 129-150 | + | – |
| *Brachypodium mexicanum* | Bmexi.01UG208900.1.p | Phytozome | 140 | 108-125 | 108-125 | + | – |
| *Camellia sinensis* | THG10281.1 | NCBI | 423 | 142-157<br>321-336 | 142-160<br>319-336 | + | – |
| *Carya illinoensis* | XP_042975722.1 | NCBI | 128 | 10-34 | 22-43 | – | – |
| *Cucurbita maxima* | XP_022995151.1 | NCBI | 178 | 7-28 | 6-26 | + | – |
| *Cucurbita pepo* subsp. *pepo* | XP_023545371.1 | NCBI | 151 | 132-150 | 132-150 | + | – |
| *Cryptomeria japonica* | XP_059077992.1 | NCBI | 144 | 96-111 | 96-114 | + | – |
| *Glycine soja* | RZB63908.1 | NCBI | 206 | 90-109 | 90-112 | + | + |
| *Gossypium barbadense* | Gobar.D06G228300.1.p | Phytozome | 159 | 137-155 | 137-155 | – | +/- |
| *Gossypium darwinii* | Godar.A06G226200.1.p | Phytozome | 159 | 138-158 | 137-157 | – | +/- |
| *Juglans regia* | XP_018844524.2 | NCBI | 205 | 20-41 | 28-47 | + | +/- |
| *Linum usitatissimum* | Lus10029623 | Phytozome | 100 | 11-31 | 12-35 | – | – |
| *Malus domestica* | XP_028945193.1 | NCBI | 205 | 43-64 | 43-68 | + | – |
| *Malus domestica* | RXH68939.1 | NCBI | 569 | 122-144<br>152-172<br>271-291 | 123-144<br>150-172<br>275-295 | – | +/- |
| *Manihot esculenta* | XP_043807697.1 | NCBI | 162 | 141-161 | 141-161 | – | – |
| *Olea europaea* subsp. *europaea* | CAA3000307.1 | NCBI | 177 | 69-96 | 57-76<br>82-99 | + | + |
| *Oryza brachyantha* | XP_040380539.1 | NCBI | 156 | 6-24 | 6-28 | + | + |
| *Oryza sativa Indica Group* | EEC71462.1 | NCBI | 328 | 11-29 | 12-33 | – | +/- |
| *Physcomitrium patens* | XP_024369173.1 | NCBI | 425 | 77-101<br>161-176 | 37-55<br>76-98<br>159-176 | + | + |
| *Picea abies* | MA_8815334g0010 | PlantGenIE | 230 | 203-225 | 206-225 | + | – |
| *Populus trichocarpa* | XP_024465836.1 | NCBI | 179 | 26-49 | 27-49 | – | – |
| *Quercus suber* | KAK7845242.1 | NCBI | 443 | 121-141<br>164-184<br>221-241 | 121-146<br>166-184<br>221-240 | + | – |
| *Salvia splendens* | XP_042031532.1 | NCBI | 236 | 12-29<br>81-98 | 12-30<br>58-75<br>82-101 | + | + |
| *Salvia splendens* | XP_042031534.1 | NCBI | 219 | 12-29<br>81-98 | 12-30<br>58-75<br>82-101 | + | + |
| *Salvia splendens* | XP_042031535.1 | NCBI | 212 | 12-29<br>81-98 | 12-30<br>58-75<br>82-101 | + | – |
| *Triticum aestivum* | XP_044417723.1 | NCBI | 204 | 43-64 | 43-67 | – | – |
| *Triticum turgidum* subsp. *durum* | VAH03188.1 | NCBI | 160 | 7-28 | 7-28 | + | +/- |
| *Vigna angularis* | XP_017425188.1 | NCBI | 198 | 27-48 | 27-48 | + | + |
| *Vigna radiata* var. *radiata* | XP_014500111.1 | NCBI | 198 | 27-48 | 27-48 | + | + |

The large diversity of proteins in the resulting set is noteworthy. They belong to 61 plant species, including representatives of Bryophyta (2 species), Gnetophyta (1 species) and Coniferophyta (1 species), monocots (11 species) and dicots (44 species). Among 95 predicted TM-HPts, 40 are obviously active as classic HPts since they have both the conserved histidine and the phosphorylation motif within the HPt domain. Additional 9 proteins can be classified as likely functional because they have a highly conserved phosphorylation site with only one aa substitution in the motif. 15 HPts harbor the conserved histidine but their phosphorylation motif is altered at more than one residue. 12 proteins, on the contrary, preserved the HPt domain with an intact phosphorylation motif except for the conserved histidine replaced by another aa. And, finally, 19 more proteins have lost both the active site and the corresponding motif (**Supplementary Table S2**).

In some plants, a single TM-HPt was found, and there were no similar proteins either within the same species or in related species of the same genus. Such case may be illustrated by *Juglans regia* genome which encodes only one predicted TM-HPt (XP_018844524.2). No other TM-HPt was found in the remaining *Juglans* genomes despite 36 species and variants of this genus are available in the NCBI genome database. At the same time, we found genera which include several species with putative TM-HPts. For example, in representatives of the *Salvia* genus several TM- HPts were found per species, up to 9 in *Salvia splendens* (**Supplementary Table S2**).

Most of the putative TM-HPts have non-transmembrane isoforms. This suggests that in the case of two or more transcript versions, mainly the canonical isoforms of the protein may be expressed. However, in some species, genes have been found that apparently encode only TM-HPt proteins (**Supplementary Table S3**). In *Aquilegia coerulea*, *Brachypodium arbuscular*, *Brachypodium mexicanum*, *Carya illinoinensis*, *Cryptomeria japonica*, *Cucurbita pepo* subsp. pepo, *Gossypium barbadense*, *Gossypium darwinii*, *Manihot esculenta*, *Malus domestica* (XP_028945193.1), *Theobroma cacao*, *Vigna angularis*, *Vigna radiata* var. radiata predicted TM-HPts are encoded by genes that have only one transcript variant. The species *Salvia splendens*, in which two genes encode only TM- containing HPts, is of particular interest. In this case, one of the genes encodes a single TM-HPt variant, while another gene encodes three protein isoforms, each with predicted TM domains. Among proteins represented in **Supplementary Table S3**, there are all variants of predicted TM-HPts with or without conserved histidine and/or phosphorylation motif.

The polypeptide length of most plant TM-HPts is relatively short, that is also typical for canonic cytosolic HPts. For example, the size of canonical *Arabidopsis* HPts which lack a TM domain is 154 ± 2 aa. The average size of proteins consisting the group under study is 200 ± 7 aa, which is rather close to the size of cytosolic HPts. The shortest (100 aa) protein found belongs to *Linum usitatissimum*. Of particular interest are most large proteins from the studied cluster that often include additional domains. Along with the HPt domain, the *Malus domestica* protein RXH68939.1 is predicted to contain a Fatty Acid Desaturase domain larger than 200 aa. The *Camellia sinensis* protein THG10281.1 is defined as an auxin-responsive protein, its HPt domain is adjacent to the AUX/IAA domain, the size of the latter also exceeding 200 aa. KAK7845242.1 of *Quercus suber* is peculiar as it is the only one in **Supplementary Table S2** that is predicted to contain two HPt domains. XP_024369173.1 of *Physcomitrium patens* can be considered the longest among the accumulated TM-HPts with a structural organization consisting only of single HPt and TM domains. Our preliminary experiments with recombinant HPts harboring inserted TM domains from natural TM-HPts showed the ability of these domains to anchor the constructed proteins to cell membranes.

The number of TM domains within TM-HPts may vary from 1 to 5. CCTOP and Phobius servers predict different numbers and/or lengths of TM domains in some proteins, but the global statistics for both algorithms are similar. The median number of TM domains per protein according to both servers is 1. The median length of the TM domain is 19 aa, and the average one is 19.3 ± 0.3 and 19.5 ± 0.2 aa according to CCTOP and Phobius, respectively. This length is a bit shorter than the average length of TM domains of cytokinin receptors (21-22 aa) [66]. TM helices are located in the proteins under study both at the N- or C-terminus, in some cases even in the middle.

It should be noted that functional phosphotransfer proteins with both TM and HPt domains in their structure are quite common in bacteria. We were able to find thousands of such proteins in the prokaryotic databases. As a rule, they differ from plant TM-HPts in their large sizes, which may exceed 1500 aa. Such sizes are often achieved due to a significant diversity of structures, which can include a number of functional domains along with HPt and TM ones [30]. For instance, the well-known TM histidine kinase RcsC of *E. coli* transfers its signal first to the membrane-bound phosphotransfer protein RcsD (890 aa) which in turn transmits it to the transcription factor RcsB [67]. Notably, among bacterial phosphotransfer proteins, there are also those very similar to plant TM-HPts both in size and structure, for example, proteins WP_083661941.1 *Actinophytocola xanthii* (168 aa) and WP_244434644.1 *Afipia* sp. P52-10 (184 aa) with predicted structures consists of one HPt and one TM domains.

### 3.2. Phylogenetic Analysis of TM Phosphotransfer Proteins

To clarify evolutionary relationships between potential TM-HPt proteins, we performed a phylogenetic analysis. It should be noted that the phylogenetic analysis of such short sequences is hardly possible due to their rather high similarity. The use of protein sequences and algorithms of the software like MEGA [68] resulted in low confidence phylogenetic trees. We carried out the analysis with MrBayes [48] using not aa but coding nucleotide sequences. This approach gave highly reliable results. Only sequences of angiosperms were used in the analysis, since adding representatives from other groups significantly complicates (and currently does not allow) obtaining a tree of sufficient quality. The list of analyzed proteins also excludes apparently inactive TM-HPts marked red in **Supplementary Table S2**. The resulting phylogenetic tree is presented in **Figure 1**.

**Figure 1.**
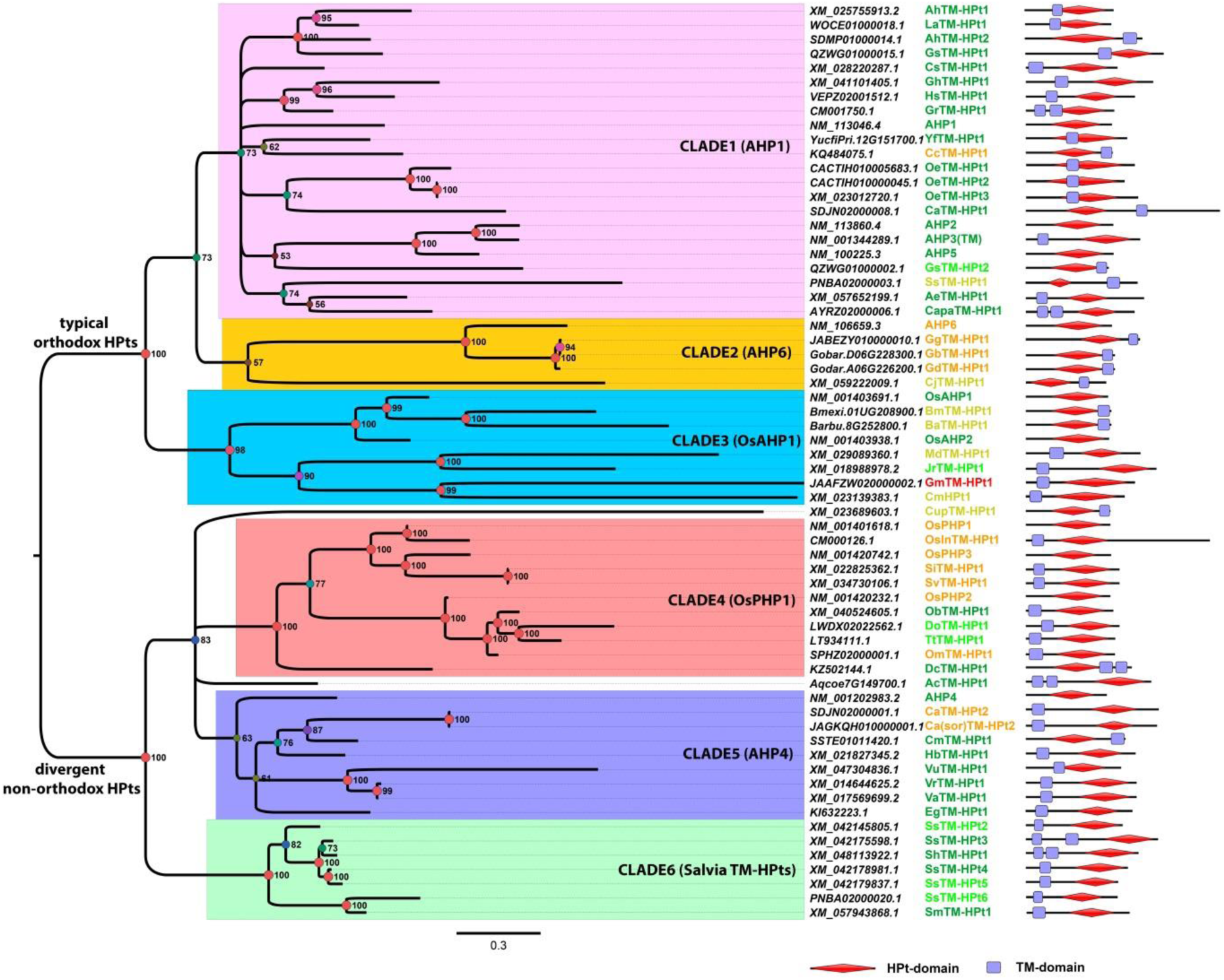
Phylogenetic tree of plant TM-HPts. The evolutionary model was the GTR substitution model with gamma-distributed rate variation across sites 6 and a proportion of invariable sites. The three columns on the right provide reference information. Among these three columns, the left one lists the transcript accession IDs. In the middle column, the color of legends provides information about the functionality of the proteins, as indicated in **Table 2** and **Supplementary Table S2**. The right column shows the domain structures of the proteins. The plant species in the figure are indicated as follows. AHP – *Arabidopsis thaliana*, OsAHP/OsPHP – *Oryza sativa* subsp. Japonica, AhTM-HPt – *Arachis hypogaea*, LaTM-HPt – *Lupinus albus* cv Amiga, GsTM- HPt – *Glycine soja* cv W05, CsTM-HPt – *Camellia sinensis*, GhTM-HPt – *Gossypium hirsutum*, HsTM-HPt – *Hibiscus syriacus* cv Baekdansim isolate YM2019G1, GrTM-HPt – *Gossypium raimondii*, YfTM-HPt – *Yucca filamentosa*, CcTM-HPt – *Cajanus cajan*, OeTM-HPt1 – *Oleae europaea*, CaTM-HPt – *Cucurbita argyrosperma* subsp. argyrosperma cv Calabaza pipiana, SsTM- HPt – *Salvia splendens*, AeTM-HPt – *Actinidia eriantha*, CapaTM-HPt – *Capsicum annuum* cv CM334, GgTM-HPt – *Gossypium gossypioides* isolate 5, GbTM-HPt – *Gossypium barbadense*, GdTM-HPt – *Gossypium darwinii*, BmTM-HPt – *Brachypodium mexicanum*, BaTM-HPt – *Brachypodium arbuscula*, MdTM-HPt – *Malus domestica*, JrTM-HPt – *Juglans regia*, GmTM-HPt – *Glycine max* cv EMBRAPABRS 537, CmTM-HPt – *Cucurbita maxima*, CupTM-HPt – *Cucurbita pepo*, OsInTM-HPt – *Oryza sativa* subsp. Indica, SiTM-HPt – *Setaria italica*, SvTM-HPt – *Setaria viridis*, ObTM-HPt – *Oryza brachyantha*, DoTM-HPt – *Dichanthelium oligosanthes* cultivar Kellogg, TtTM-HPt – *Triticum turgidum* subsp. Durum, OmTM-HPt – *Oryza meyeriana*, DcTM-HPt – *Dendrobium catenatum*, AcTM-HPt – *Aquilegia coerulea*, Ca(sor)TM-HPt – *Cucurbita argyrosperma* subsp. sororia isolate JBR-2021, CmTM-HPt – *Cucumis melo* var. makuwa cv SW 3, HbTM-HPt – *Hevea brasiliensis*, VuTM-HPt – *Vigna umbellata*, VrTM-HPt – *Vigna radiata*, VaTM-HPt – *Vigna angularis*, EgTM-HPt – *Mimulus guttatus* cv IM62/*Erythranthe guttata*, SmTM-HPt – *Salvia miltiorrhiza*, ShTM-HPt – *Salvia hispanica*.

The overall tree topology was similar to that generated previously [69]. As in the mentioned work, we constructed a tree only for representatives of angiosperms due to the difficulties in establishing the exact position in the tree of the branches of non-angiosperm HPts. Each transcript/protein (**Supplementary Table S4)** represented in the tree (**Figure 1**) corresponds to a distinct gene. Our tree has two large branches, which we designated as typical orthodox HPts and divergent non-orthodox HPts. According to the reference species *Arabidopsis thaliana* and *Oryza sativa* sub-species Japonica, the first group includes orthologs of AHP1-3,5,6 and OsAHP1 and 2, while the second group includes orthologs of AHP4 and OsPHP1-3. It should be noted that the Arabidopsis HPt AHP4 stands apart, as it has small structural and significant functional differences from other HPts of this plant; in particular, it hardly supports canonical phosphotransfer into the nucleus [70]. Each branch in turn is divided into three clades. We named each clade after a typical representative or, in the case of clade 6, according to a particular plant genus (*Salvia*). Clades 1 and 2 do not contain representatives of the grasses from Monocots. Clade 2, however, consists only of PHPs – orthologs of phosphorelay inhibitor AHP6 from Arabidopsis. Among members of Clade 3, there is only one potentially active HPt of *Juglans regia*. Most likely, dicots tend to reduce genes belonging to Clade 3, as indicated by the longest branch length (accelerated mutagenesis) in the entire tree for the remaining three dicots in this clade, *Malus domestica*, *Glycine max*, and *Cucurbita maxima*. Clades 4 and 5 include representatives of monocots and dicots, respectively. In the fourth clade, not all proteins are PHPs, as is typical for *Oryza sativa* sub-species Japonica. The *Oryza brachyantha* member is an apparently active HPt. Similarly, clade 5 contains both active and pseudo HPts.

Notably, HPts of the genus *Salvia* were allocated in a separate clade, which has not been recognized before. For example, in the work [69], where the phylogeny of cytokinin signaling components was considered in detail, this clade was absent, since the sample did not include the corresponding plant species. According to our preliminary data, this clade is present in potato, tomato, *Hevea brasiliensis* and a number of other plants, but is still absent in most species. In general, it can be assumed that the TM domain is found in all known phylogenetic clades in angiosperms, most often single and upstream of the HPt domain. At the same time, no reliable relationship was found between the phylogenetic status of the TM-HPt and the number and location of TM domains in the protein. Apparent distinctive traits, such as those in clades 2 and 6, are preferably explained by the close relationship of the plant species involved. This means that in this case the most likely scenario is that this gene series originated from a single TM-HPt gene in an evolutionarily close ancestor of the group at the origin of the genus or at the formation of subgenera within the genus.

The emergence of TM-HPt lines at the stage of genera or subgenera formation is observed in several cases, when species of one genus form clusters, in which species from other genera are absent. This is observed in the genera: *Salvia* (Clade 6), *Vigna* (Clade 5), *Brachypodium* (Clade 3), *Setaria* (Clade 4), *Gossypium* (Clade 2). It is interesting that in the genus *Gossypium* TM-HPts appeared not only in clade 2 (GgTM-HPt1, GbTM-HPt1, GdTM-HPt1), but also in clade 1 (GhTM-HPt1, GrTM-HPt1). But a representative from *Hibiscus sinensis* “wedges” itself into the latter clade. In several species, TM-HPts seem to independently arise several times. In *Arachis hypogea* – in Clade 1 (AhTM-HPt1 and 2), in *Salvia splendens* – in Clade 1 (SsTM-HPt1) and many in its own Clade 6 (SsTM-HPt2-6), in *Glycine soja* – in Clade 1 (GsTM-HPt1 and 2). Thus, we can conclude about active and largely independent evolution of phosphotransfer proteins towards the formation of TM-HPts in angiosperms, and, taking into account the non-angiosperm species not included in the tree, in all land plants as a whole.

### 3.3. Transcriptomic Studies of Transmembrane HPt genes

The question whether transcripts encoding TM-HPt proteins are expressed was addressed using a transcriptomic approach in 12 plant species covering various classes of TM-HPt proteins grouped in **Supplementary Table S1.** The analysis revealed that most of the investigated transcripts encoding functional TM-HPt proteins are more rare in comparison to two frequently used reference genes (**Figure 2**). The highest expression was found in the transcript of the *Physcomitrium* TM-HPt, which reached a moderate transcription level comparable to the PP2AA2 reference transcript, closely followed by the transcript encoding the *Camellia sinensis* protein THG1028.1. Of 8 functional TM-HPt transcripts investigated (green color), the one of *Hevea brasiliensis* can be regarded as absent. Two of the 3 annotated transcripts encoding non-functional TM-HPt proteins (red color) are virtually not transcribed too. The transcript of *Cucurbita pepo*, whose encoded protein contains an active phosphorylation site but a significantly altered motif (yellow bar) was at best marginally transcribed.

**Figure 2.**
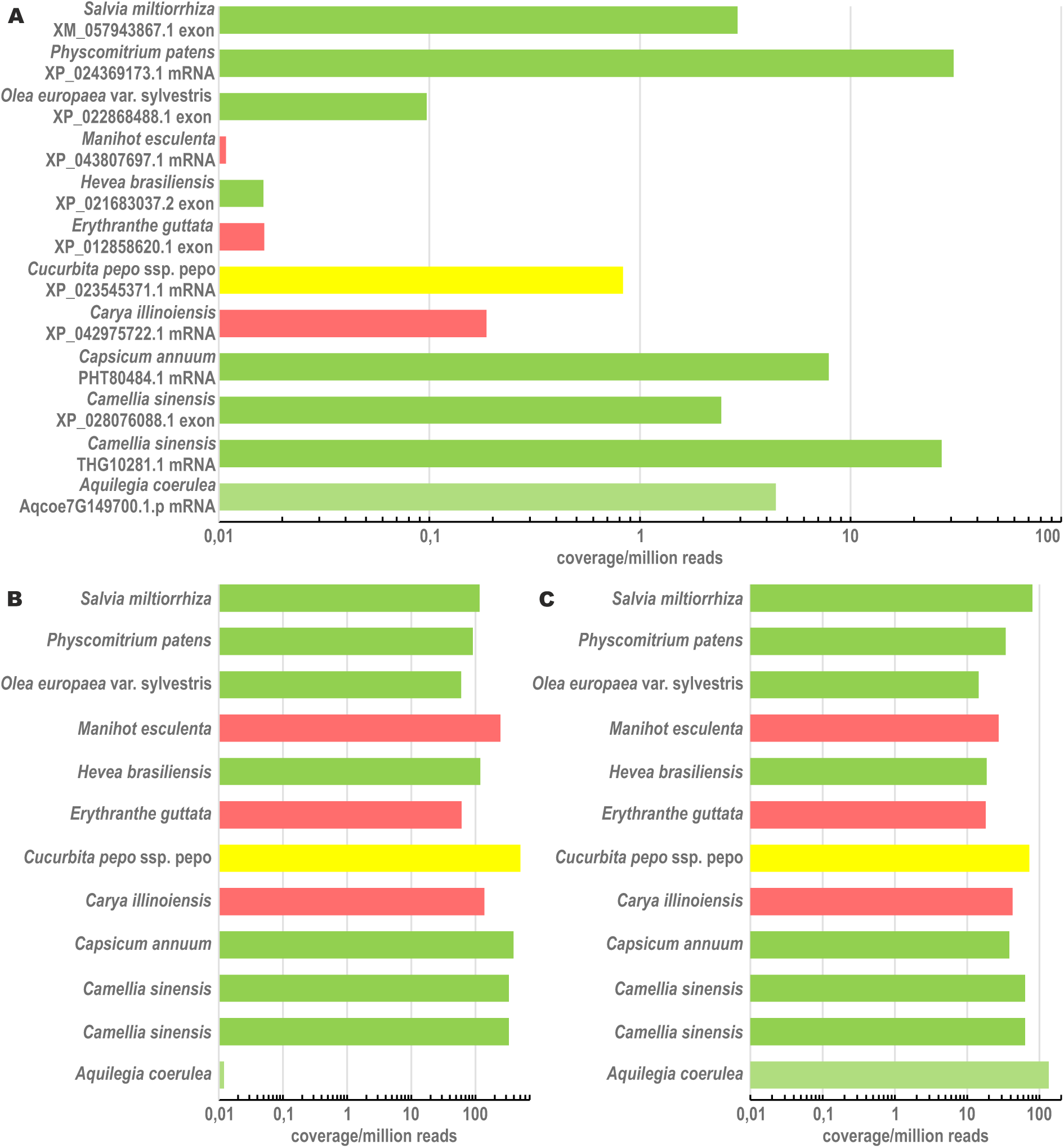
Expression of the transcripts encoding TM-HPt proteins (A) and of the orthologs of the two *Arabidopsis thaliana* reference genes *UBC10* (AT5G53300, B) and *PP2AA2* (AT3G25800, C). Expression of the transcripts is estimated by the maximum coverage, plotted as coverage per million reads. For each TM-HPt transcript, the respective protein ID is given. In case of a single transcript, the entire mRNA was used for maximum coverage calculation (identified as “exon”). The color-code of the bars in (A) is the same as in **Table 2** and **Supplementary Tables S2** and **S3**. For the reader convenience, the color of the bars corresponding to reference genes is the same as in (A).

In summary, it can be stated that transcripts encoding functional TM-HPt proteins are expressed, but most of them at rather low levels. The highest transcription rate (maximum coverages between 1 and 100 per million reads) was found in transcripts encoding proteins with a fully functional phosphorylation motif (green bars). Intriguingly, two of the three annotated transcripts encoding non-functional TM-HPts (red bars), are virtually not expressed. These non-functional genes may have become pseudogenes. The only “red” gene (from *Carya illinoiensis*), that appears to be marginally expressed, encodes a protein completely lacking the conserved phosphorylation motif. This severely mutated protein may no longer take part in MSP signaling. The rather high expression level of the *Physcomitrium* TM-HPt transcript may indicate a more prominent role of TM-HPts in early land plants; however, more TM-HPt transcripts of mosses and pteridophytes would have to be studied to corroborate this notion.

In fact, the moderate-to-low transcript content of many TM-HPt genes in the whole plant organism may be indicative of the organ- and/or tissue specificity of these genes. It also cannot be ruled out that the expression of this group of genes may be confined to a specific stage of plant development or controlled by inducible promoters. Further detailed studies of the genomes and transcriptomes of the above-mentioned plant species will clarify this issue.

### 3.4. Molecular Modeling of Transmembrane HPts

To evaluate the possible functionality in more detail as well as to investigate the spatial organization of the studied proteins, structural models of different TM-HPts were built. Their orientation in the membrane was predicted using the PPM 3.0 service [62,63]. Some of these models were optimized for positioning in the membrane profile since they were initially constructed without appropriate constraints.

We used the canonical soluble HPt 3D shape (AHP 1–3, 5) as a reference when comparing the resulting TM- HPts models. The canonical structure of HPts involved in MSP signaling consists of 6 α-helices. Three out of the six α-helices (α2, α3 and α4) are involved in the formation of the interaction interface with the receiver domain (RD) of the receptor kinase [71]. The protein-protein interaction interfaces of HPt homodimer and HPt–HKRD heterodimer, significantly overlap [15].

The diversity of topology and folding of the obtained models of different TM-HPts was in the number of TM domains, the completeness of the canonical HPt part, the spatial arrangement features and size of the extracytosolic part. We use the term «extracytosolic» for protein regions located on the side of the membrane opposite to the canonical HPt moiety. In this case, it does not matter on which side of the membrane the prediction software placed this fragment. This is due to the fact that classical HPts have been shown to preferentially localize in the cytosol and the nucleus, but not in the apoplast [72,73]. We distributed all the resulting models into several groups (**Figure 3**).

**Figure 3.**
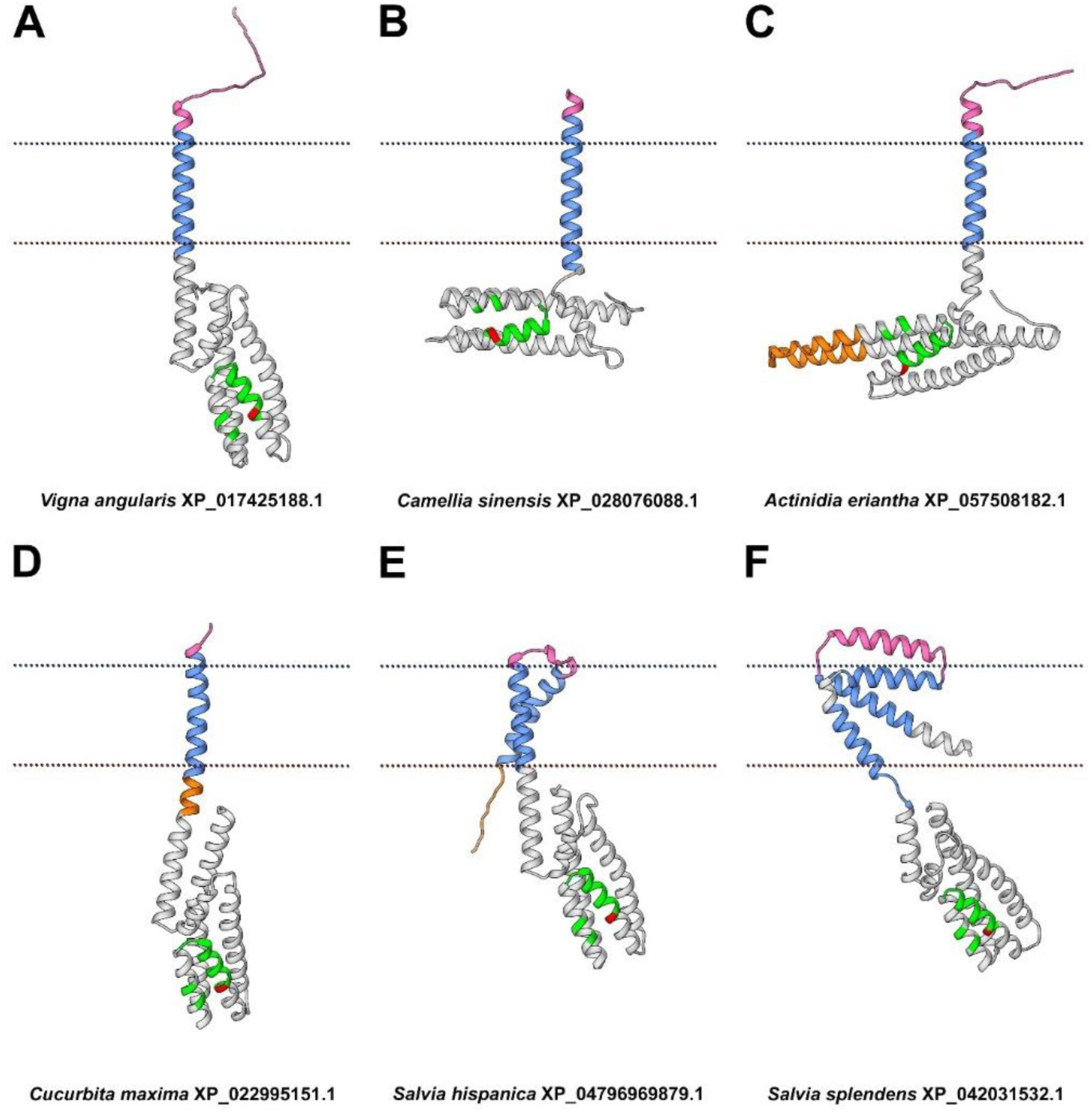
Selected molecular models of TM-HPts categorized by topology and folding features. Each model represents one of the structural groups to which it has been categorized. **(A)** *Vigna angularis* XP_017425188.1 belongs to a group 1; **(B)** *Camellia sinensis* XP_028076088.1 – group 2; **(C)** *Actinidia eriantha* XP_057508182.1 – group 3; **(D)** *Cucurbita maxima* XP_022995151.1. – group 4; **(E)** *Salvia hispanica* XP_047969879.1 – group 5; **(F)** *Salvia splendens* XP_042031532.1 – group 6; The TM segment is highlighted in blue, the cytosolic insertion is yellow, the extracytosolic region in pink. The conserved motif is shown in green, the phosphoaccepting histidine in red.

In proteins classified to the first group, the basic HPt part is mostly preserved, the TM domain is a direct elongation of the N-terminal α1-helix of the HPt moiety and the extracytosolic fragment is mostly unstructured (**Figure 3A**). Such proteins are *Vigna radiata* XP_014500111.1 and *Vigna angularis* XP_017425188.1. In both *Vigna* spp. proteins, the canonical HPt part is preserved completely, extracytosolic part is short, mostly unfolded and includes an N-glyco motif (according to Protter). The N-glyco motif is the site of N-glycosylation, the biochemical process of attaching a glycan to the *N*4 atom of asparagine residue. N-glycosylation has a structural function, affects protein stability and solubility, protects proteins from aggregation, and can mediate signal transduction in cells [74,75].

The second group of the HPt set includes proteins whose α1-helix of the canonical HPt part is significantly shortened and continuously transitioned to TM or completely replaced by TM helix (**Figure 3B**). *Camellia sinensis* XP_028076088.1, *Populus trichocarpa* XP_024465836.1, *Malus domestica* XP_028945193.1, and *Actinidia eriantha* XP_057508184.1 belong to this type. The extracytosolic part of this protein group could be different. Thus, in *Camellia sinensis* XP_028076088.1 it is a short continuation of the TM helix (5 aa in total). In *Populus trichocarpa* XP_024465836.1 it is a long α-helix (26 aa). In *Malus domestica* XP_028945193.1 there is a large unfolded region 42 residues long containing an N-glyco motif.

*Actinidia eriantha* XP_057508182.1 (**Figure 3C**) is referred to third structural group. Together with *Olea europaea* XP_022868488.1, these two proteins have an insertion within the canonical HPt part itself. *Actinidia eriantha* XP_057508182.1 has an insertion caused by the elongation of α5 and α6 helices of the HPt fragment. In *Olea europaea* XP_022868488_1, atypically arranged TM regions localize not at the termini of the canonical HPt domain, but in the middle of it as an insertion between α3 and α4 helices.

In the fourth structural group (**Figure 3D**), elongation rather than shortening of the cytosolic portion of the α1-helix is observed, i.e. there is an additional fragment between the canonical HPt part and the TM region. The model of *Cucurbita maxima* XP_02299995151.1 have such elongation. In *Cucurbita maxima* XP_02299995151.1, the N-terminal region, the TM domain and the elongated α1 of canonical part represent a single continuous α-helix. Again, the HPt domain here contains an N-glyco motif.

The TM-HPts of *Salvia spp.* are of particular interest in terms of structural diversity. Their HPt proteins include from one to three TM domains. The considered *Salvia hispanica* proteins have two N-terminal TM domains connected by a short linker (**Figure 3E**) and belong to the fifth structural group. The difference between XP_047969881.1 and XP_047969879.1 is in 17 aa shortening of the C-terminal α6-helix of the canonical HPt part of the first protein.

Nine TM-HPts were found in *Salvia splendens*, three of them have three TM domains and six others have one TM domain each. The spatial shape of all three proteins with three TM helices (XP_042031532.1, XP_042031534.1, and XP_042031535.1) is similar, which allowed us to put them into a sixth group (**Figure 3F**). Between the first and the second TM motifs there is a helix-shaped insertion comparable in size to the TM segments. The third TM domain is connected to the canonical HPt part flexibly via a loop. These proteins are distinguished by the length of the C-terminal α6-helix of the canonical part. In XP_042031532.1 this part is full-sized, whereas in XP_042031534.1 the C-terminal α-helix is reduced by about a half, and in XP_042031535.1 the canonical part comprises only five α-helices.

Among the obtained models which are not considered in detail in this paper, several other groups can be additionally distinguished. First, these are TM-HPts with a signal sequence in the extracytosolic part. These include XP_025611698.1 of *Arachis hypogaea* and XP_047160793.1 of *Vigna umbellata*. Second, these are phosphotransfer proteins with a single TM domain, but located at the C-terminus rather than at the N-terminus. Such HPts may be exemplified by those of *Picea abies* MA_8815334g0010 and *Gossypium darwinii* Godar.A06G226200.1.p. Finally, there are proteins that have two predicted potentially functional domains. *Quercus suber* KAK7845242.1, according to the model, includes two predicted HPt domains separated by three TM domains, a small extracytosolic region, and several cytosolic domain extensions. *Camellia sinensis* THG10281.1 has a canonical HPt domain with all six preserved α-helices located at the N-terminus, as well as PB1 domain typical for auxin signaling proteins, at the C-terminus.

### 3.5. Possible Functional Meaning for the Presence of the TM Domain in Plant HPts

The functional meaning of the TM domains in plant HPts obviously depends primarily on the structural features of the TM-HPt itself. As this work has shown, the ability of TM-HPts to specifically transfer a “hot” phosphate varies over a wide range, where at one pole are proteins with preserved canonical HPt domain and an impeccable phosphorylation site, while at the opposite pole are proteins with heavy violations of both. It is natural to assume that the functional role of proteins with opposite structural features will most likely also be different. Membrane proteins with a conserved histidine capable of transferring signal in the form of mobile phosphate can create branches from the canonical MSP signaling to the nucleus, directing it to certain membrane-bound targets. This version is supported by our preliminary data on the presence in plants of not only membrane HPts, but also membrane RRs. Moreover, the suggested structural diversity of TM-HPts argues for putative multivariate targets in the cell beyond the genetic material. As an analogy, we can refer to bacterial HPts, many of which are membranous proteins with complex multidomain structures [30].

Theoretically, it is also possible to transfer a phosphate from TM-HPt to a canonical soluble RRB. Although the majority of RRBs are believed to be localized in the nucleus, it cannot be ruled out that some of their representatives may partially reside in the cytoplasm and be able to directly interact with TM-HPts. This may result in a targeted interaction of TM-HPt with a specific RRB(s), similar to the specific interaction of components of a two-component system in bacteria [11], which in turn may result in a change in the expression of a more specialized set of genes. Another possibility for the noticeable influence of TM-HPts on the action of the MSP system is that these proteins may form a kind of low-active pool on membranes. Such a pool can be, in principle, capable of rapidly switching to an active state in the case of cleavage of the polypeptide chain at some point between TM and HPt domains. Such cleavage is possible upon induction of the expression and/or activity of a specialized endoprotease. Considering the huge number of protease genes identified in plants (>650 and >800 in rice and Arabidopsis, respectively, [76], such a scenario seems quite plausible. Moreover, the sign of the effect – positive or negative – would directly depend on the main characteristic of the cleaved TM-HPt. In the case of cleavage of active proteins (green in **Table 2**, **Supplementary Tables S2** and **S3**, and **Figures 1** and **2**), MSP should be enhanced, and in the case of cleavage of phosphotransfer inhibitors lacking a conserved histidine (orange in **Table 2**, **Supplementary Tables S2** and **S3**, and **Figure 1** and **2**), it should be correspondingly weakened.

Another option for the negative effect of TM-HPt on phosphorelay signaling may be the preserved ability of membrane-bound HPt to dimerize. As a result, not only TM-HPt itself will lose its ability to transmit a signal to the nucleus, but it also will bind mobile HPts, reducing their active concentration in the cell and thereby weakening the canonical MSP. Thus, TM-HPts can act as modulators of the strength and specificity of plant sensory histidine kinase signaling, including directing MSP signaling to new and yet unknown, non-canonical targets.

In general, the accomplished work has opened up new, previously unknown areas of science that can lead to a breakthrough in our knowledge of the molecular mechanisms of action of phytohormones (cytokinins) and MSP-based plant signaling systems.

## 4. Conclusion

A global search for a special class of phosphotransfer proteins harboring TM aa segment(s) and belonging to MSP system of plants has been performed using large-scale bioinformatics methods. The analysis of more than 120 sequenced plant genomes revealed about a hundred predicted phosphotransfer proteins with the unconditional presence of TM domains, identified by rigorous independent algorithms (**Table 2, Supplementary Table S2**). Totally, about a hundred of such proteins from 61 species were uncovered, most of them with preserved functional domains and thus considered potentially active. Phylogenetic analysis has divided these TM-HPts into two main groups, one representing orthologs of Arabidopsis’ AHP4, while another – orthologs of all other rather uniform HPts (AHP1–3, 5) of this species. Although most of these proteins are isoforms of canonical soluble HPts encoded by distinct transcript versions, about two dozen genes in 13 plant species were found expressing a single mRNA version encoding TM protein (**Supplementary Table S3**). Together with the transcriptomic data that most of TM-HPt encoding transcripts are surely formed in the cell *in vivo* (**Figure 2**), this leaves virtually no doubt about the real existence of such proteins in the cells of many plants. The structural models of the revealed proteins were built using molecular modeling methods and a wide variety of model structures have been demonstrated. The analysis showed the potential of TM- HPts to both positively and negatively influence the MSP signaling. New, as yet unexplored signaling pathways also cannot be ruled out. This work laid the foundation for a targeted study of non-canonical membrane branches of the MSP pathway in many plant species. Among the latter, there are such economically valuable species as rice, wheat, soybean, sunflower, cotton, etc. This indicates the particular importance of this newly discovered area of research not only for fundamental science but also for its practical application.

## Author Contributions

Conceptualization, L.S.N., E.M.S., G.A.R.; funding acquisition, E.M.S; investigation, L.S.N., E.M.S., D.V.A.,

W.G.B.; visualization, D.V.A., W.G.B.; writing – original draft, L.S.N., E.M.S., D.V.A., W.G.B.; writing – review & editing, G.A.R.

## Acknowledgements

We are especially grateful to Petrenko Alexey Eduardovich for the creation of a special algorithm allowing us to perform GeneBank-wide screening for predicted transmembrane phosphotransmitters. We are grateful to the Ministry of Science and Higher Education of the Russian Federation for its assistance (theme No. 122042700043-9) to maintain Institute building and facilities in proper condition.

## Supplementary Materials

Supplementary materials contain:

**Table S1.** RNA-Sequencing datasets used for the transcriptomic approach to determine expression of TM-HPt-encoding transcripts.

| Species | Protein ID | Dataset (GEO ID) |
| --- | --- | --- |
| *Aquilegia coerulea* | Aqcoe7G149700.1.p | GSE158507 |
| *Camellia sinensis* | XP_028076088.1 | GSE198198 |
| *Camellia sinensis* | THG1028.1 | GSE198198 |
| *Capsicum annuum* | PHT80484.1 | GSE240948 |
| *Carya illinoensis* | XP_042975722.1 | GSE179336 |
| *Cucurbita pepo* ssp. *pepo* | XP_023545371.1 | GSE205063 |
| *Erythranthe guttata* | XP_012858620.1 | GSE280929 |
| *Hevea brasiliensis* | XP_021683037.2 | GSE101568 |
| *Hibiscus syriacus* | KAE8670382.1 | GSE99329 |
| *Olea europaea* | XP_022868488.1 | GSE145100 |
| *Physcomitrium patens* | XP_024369173.1 | GSE138467 |
| *Salvia miltiorrhiza* | XP_057799853.1 | GSE100970 |

**Table S2.**
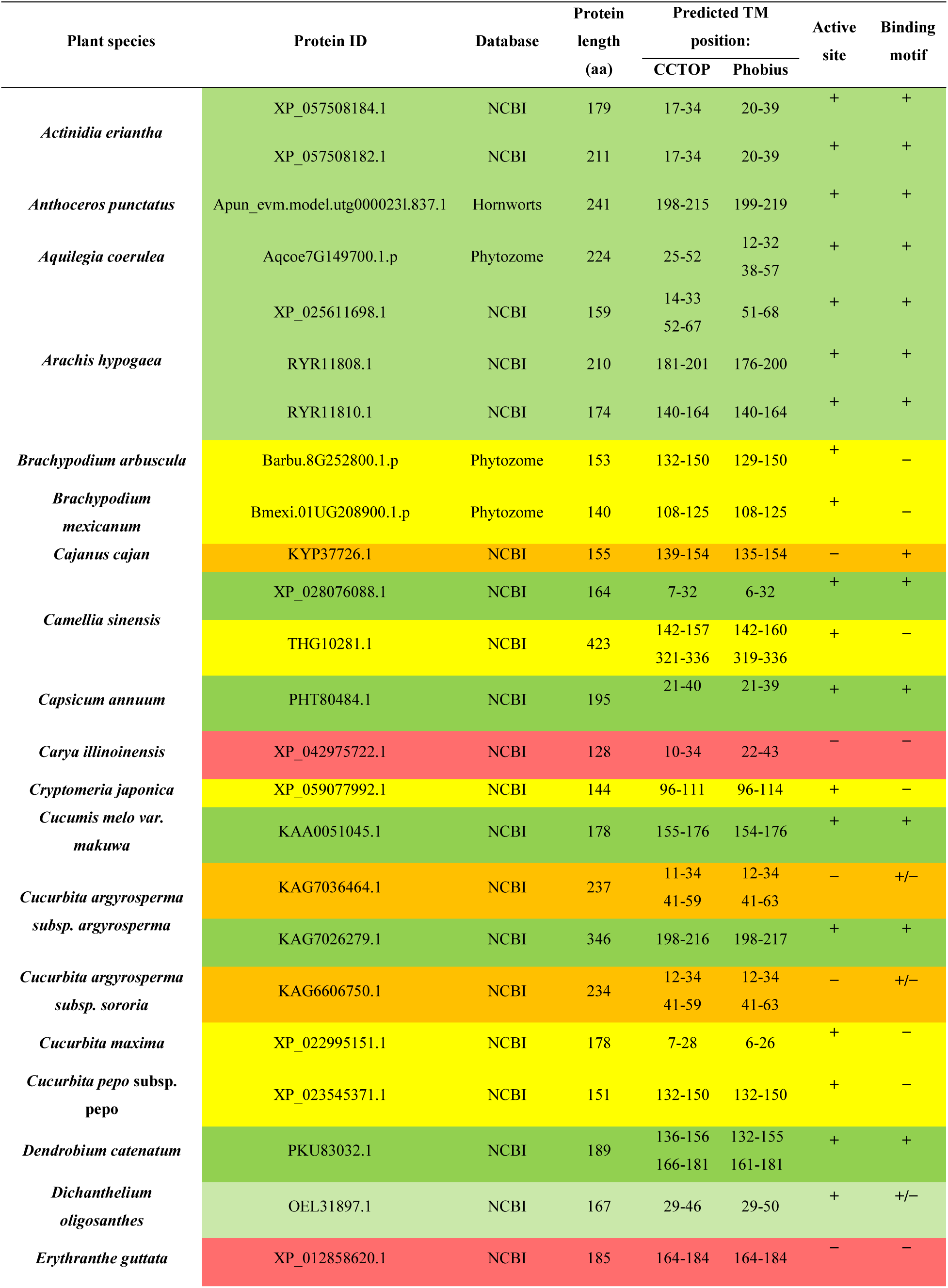
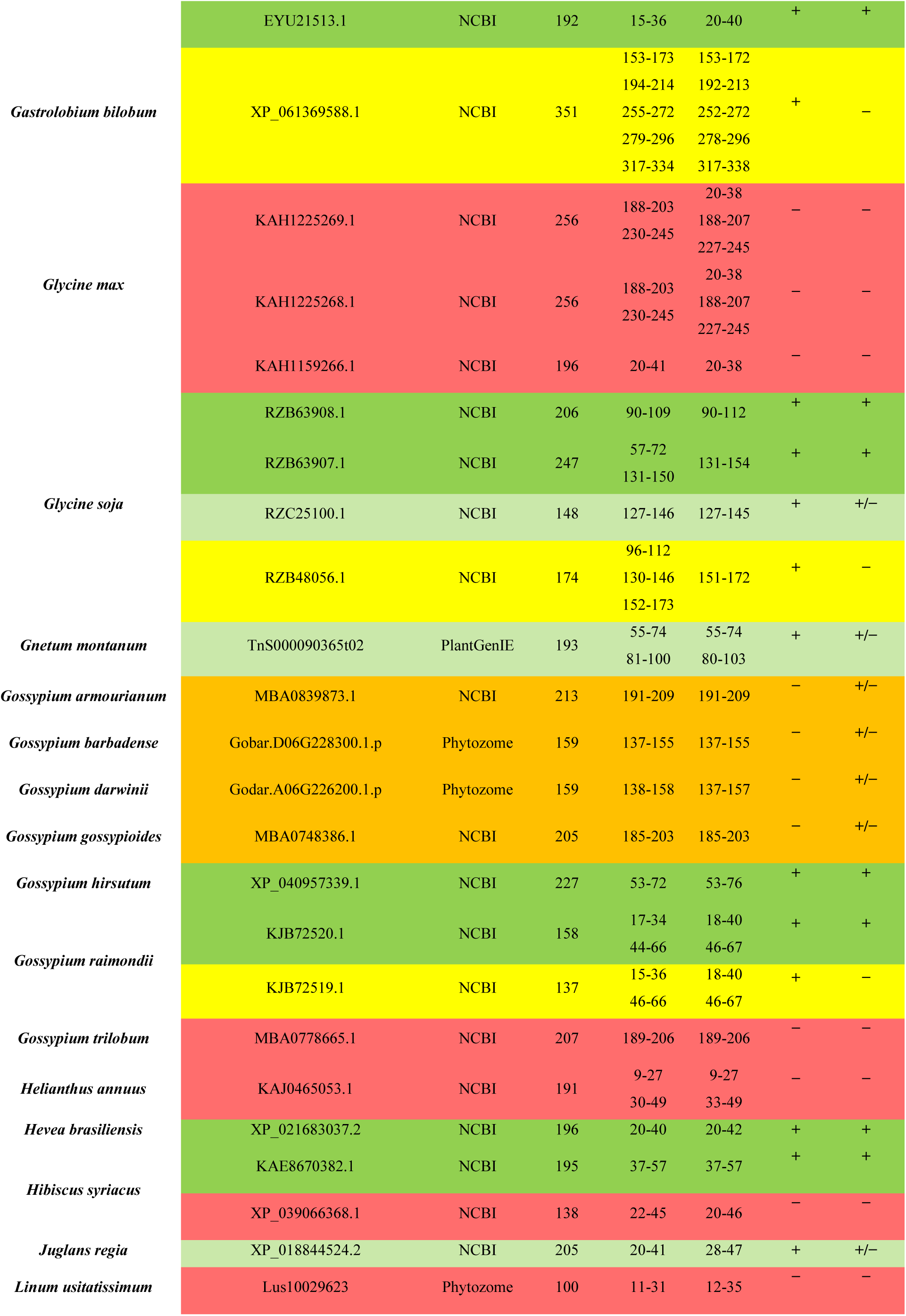
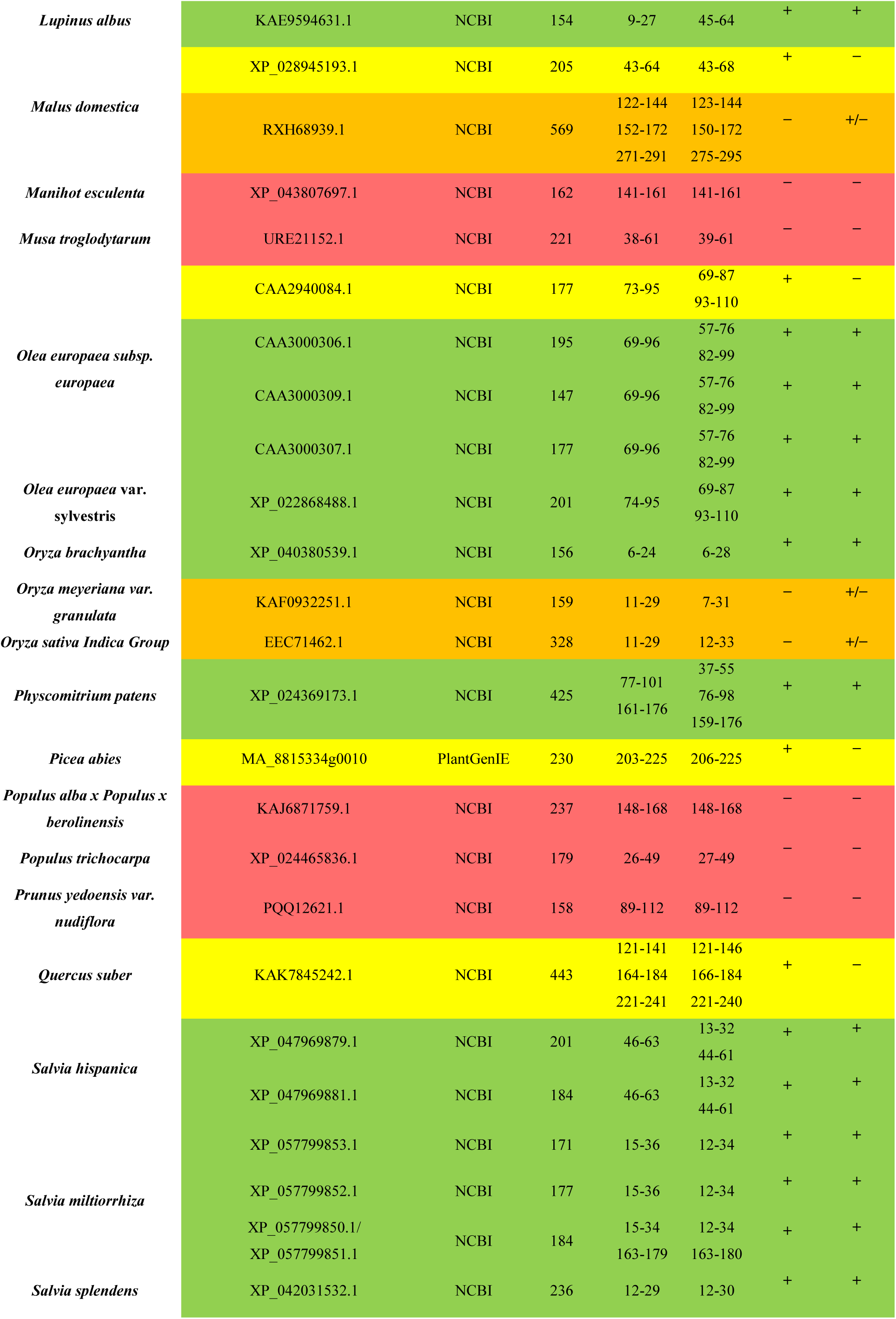
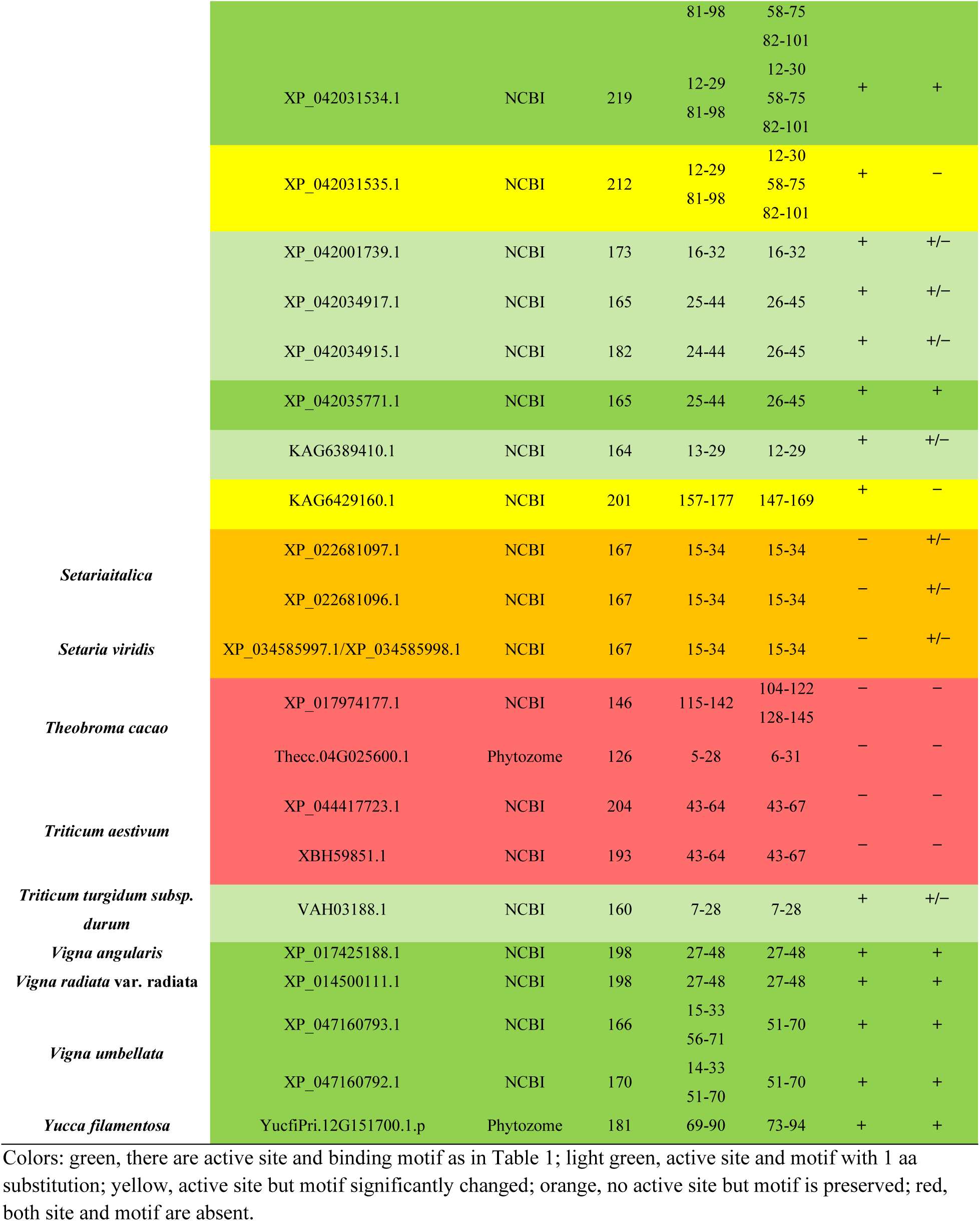
List of potential TM-HPts and their main characteristics.

**Table S3.**
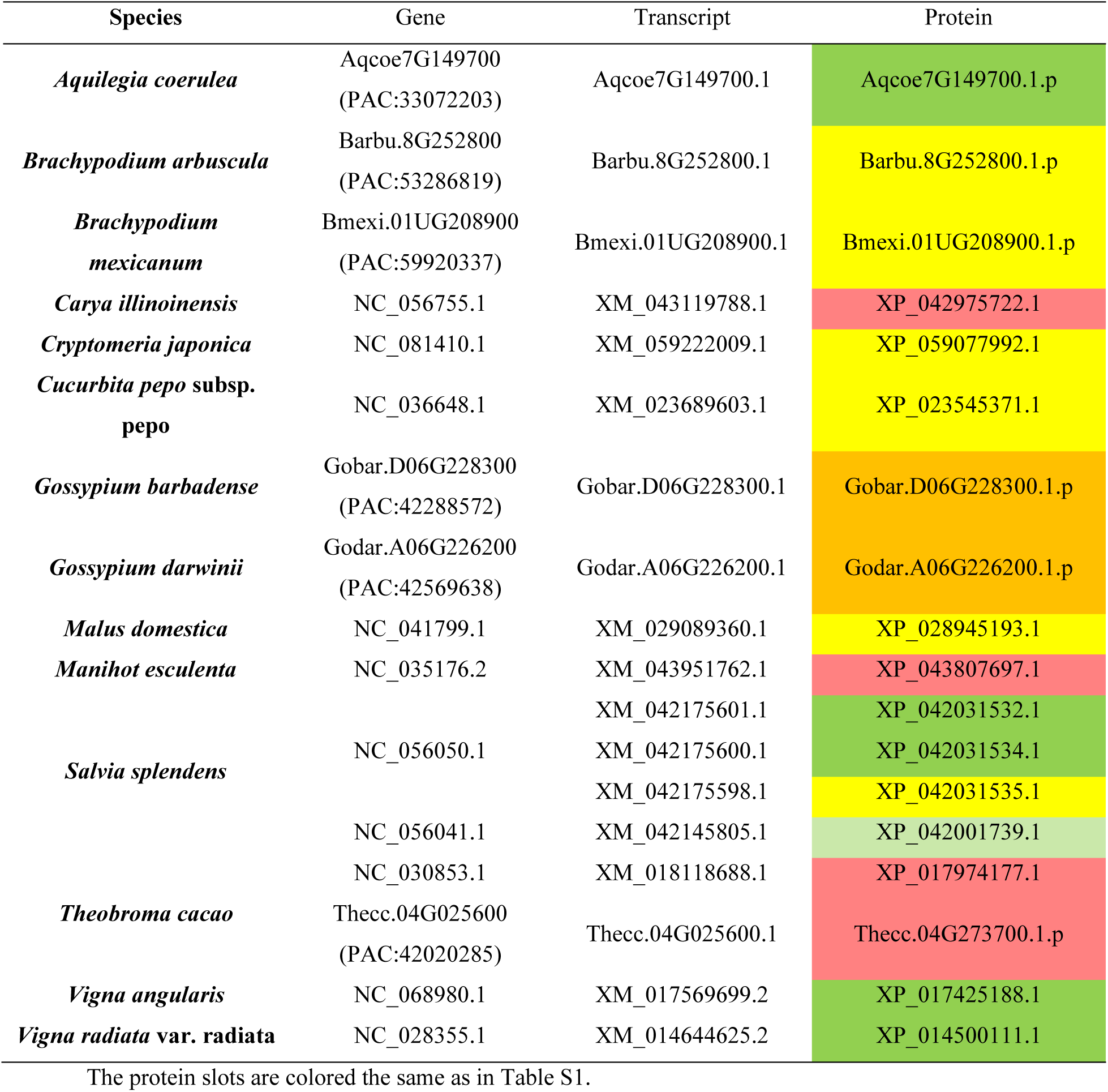
List of plant species with genes encoding only HPts with predicted TM domains.

**Table S4.**
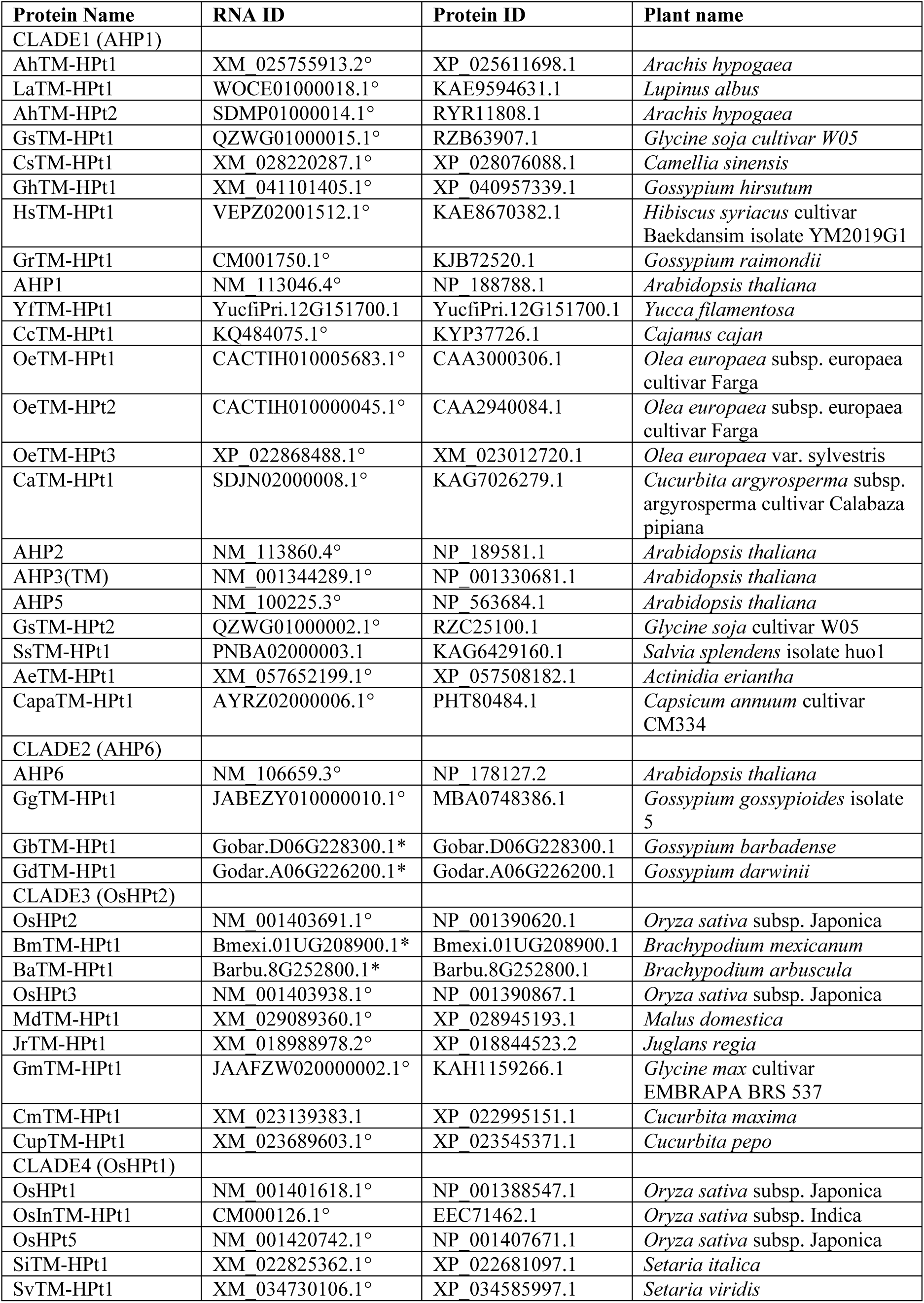
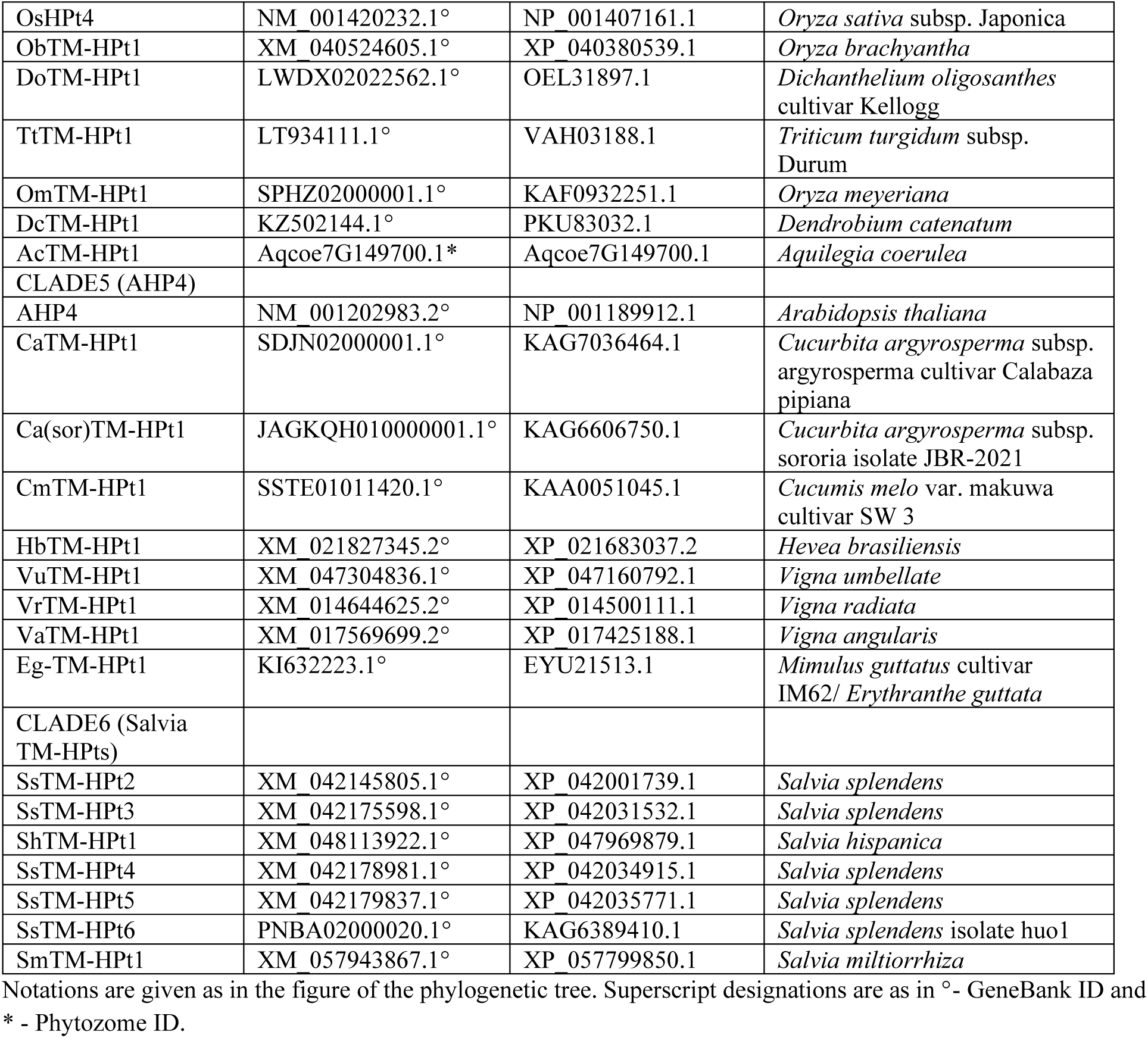
Correspondence of protein and CDS/RNA sequences used to generate the phylogenetic tree.

**Figure S1.**
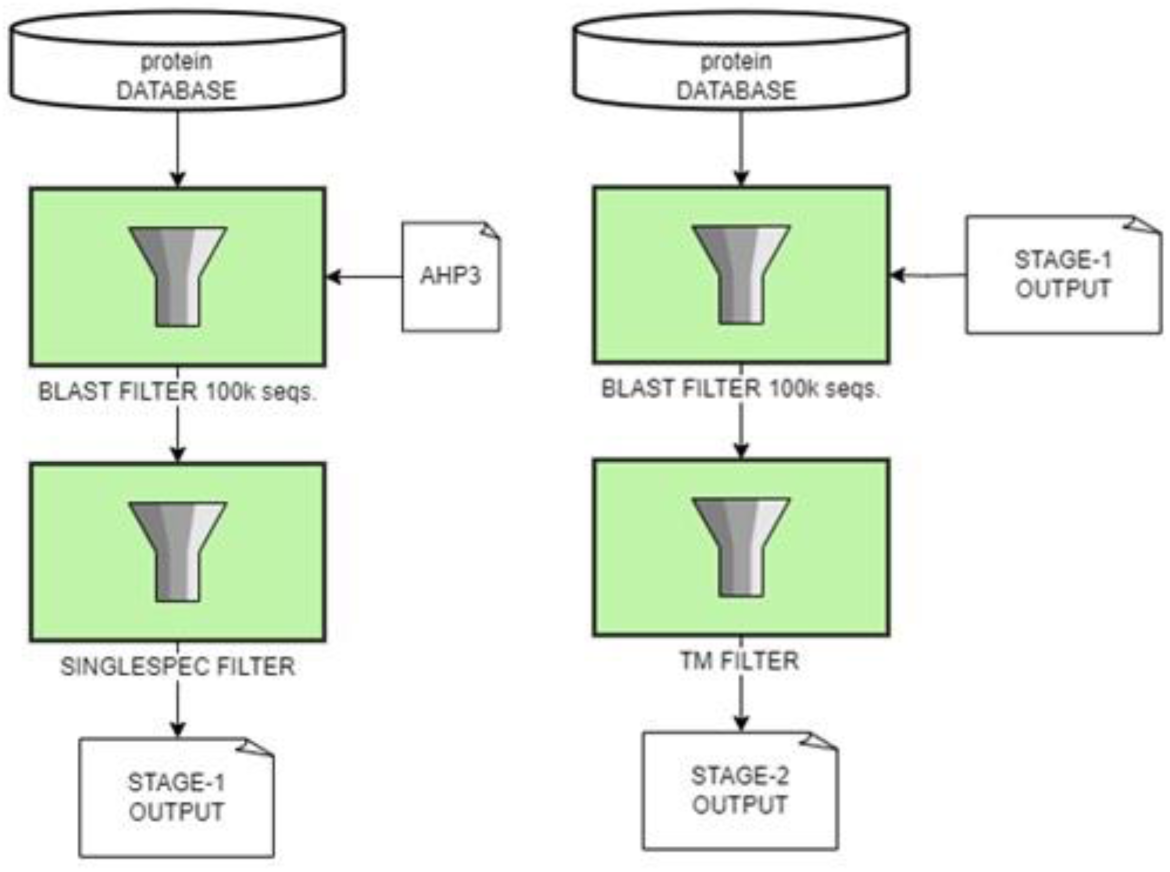
Algorithm of the automatic search for TM-HPts in the NCBI database.

**Figure S2.**
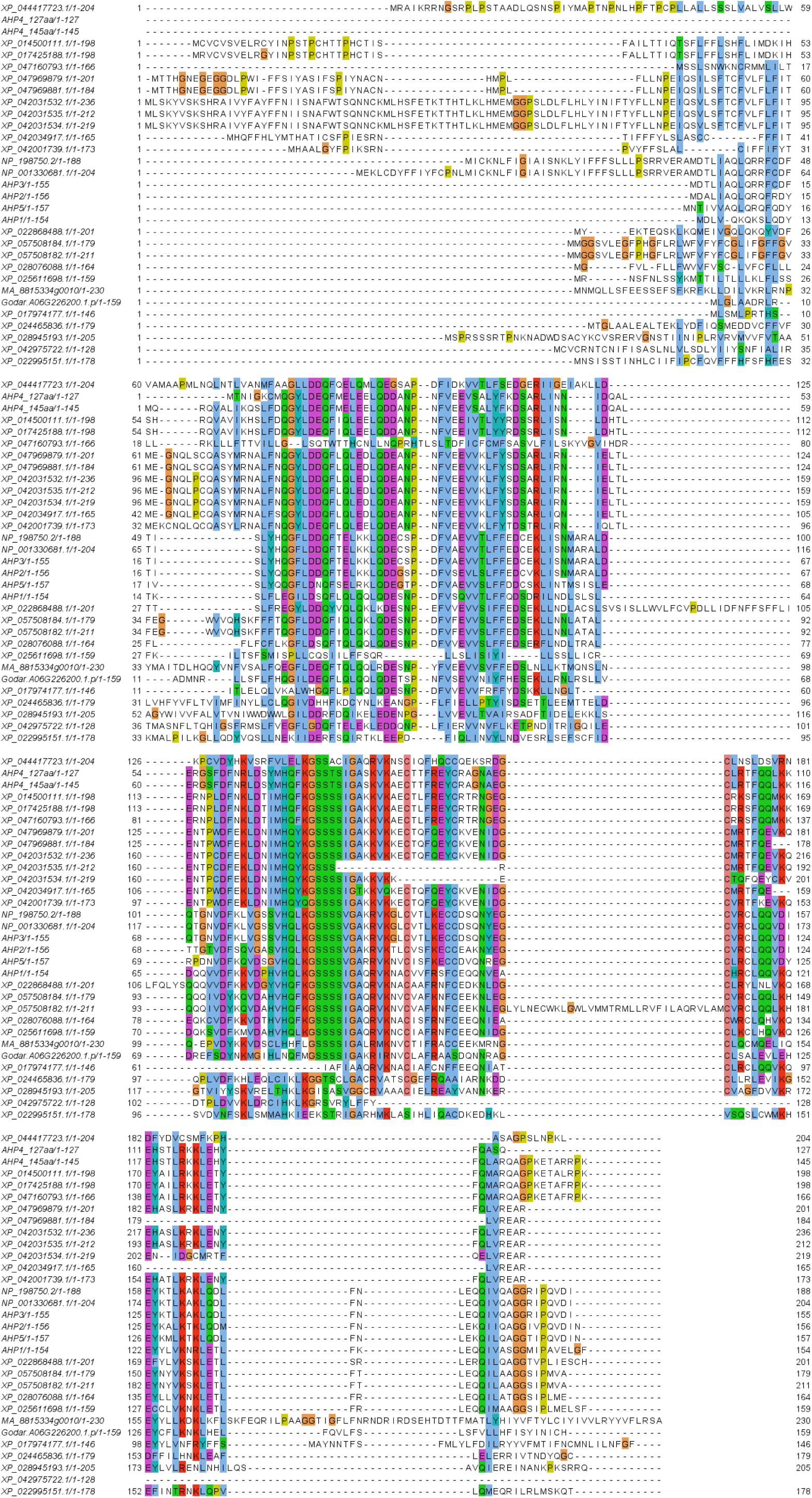
Example for TM-HPt aa sequences alignment, conserved motifs are clearly shown by vertical color bars. Protein ID numbers are given on the left.

